# AlphaFold2 transmembrane protein structure prediction shines

**DOI:** 10.1101/2021.08.21.457196

**Authors:** Tamás Hegedűs, Markus Geisler, Gergely Lukács, Bianka Farkas

## Abstract

Transmembrane (TM) proteins are major drug targets, indicated by the high percentage of prescription drugs acting on them. For a rational drug design and an understanding of mutational effects on protein function, structural data at atomic resolution are required. However, hydrophobic TM proteins often resist experimental structure determination and in spite of the increasing number of cryo-EM structures, the available TM folds are still limited in the Protein Data Bank. Recently, the DeepMind’s AlphaFold2 machine learning method greatly expanded the structural coverage of sequences, with high accuracy. Since the employed algorithm did not take specific properties of TM proteins into account, the validity of the generated TM structures should be assessed. Therefore, we investigated the quality of structures at genome scales, at the level of ABC protein superfamily folds, and also in specific individual cases. We tested template-free structure prediction also with a new TM fold, dimer modeling, and stability in molecular dynamics simulations. Our results strongly suggest that AlphaFold2 performs astoundingly well in the case of TM proteins and that its neural network is not overfitted. We conclude that a careful application of its structural models will advance TM protein associated studies at an unexpected level.

## Introduction

Although enormous resources were devoted to predict proteins’ structure for many decades, predicting a protein structure from its sequence remained a challenging task^1^. There was a change in the last Critical Assessment of Protein Structure Prediction (CASP) competition^2^ when two neural network based approaches, RoseTTAFold^3^ and AlphaFold2^4^ (AF2), were excelled. Importantly, DeepMind generated AF2-predicted structures for the human^5^ and 20 other proteomes and they were deposited to EBI (https://alphafold.ebi.ac.uk). Moreover, to ease the running of predictions for researchers, DeepMind^6^ and community Google Collaboration notebooks^7^ have been generated, albeit applying some simplifications. AlphaFold2 was trained using multiple sequence alignments (MSA) and experimental protein structures deposited before April 2018. Five different models were trained (e.g. with different random seeds, with or without structural templates) to promote an increased diversity in structure predictions^6^. The input for prediction is the sequence of a single protein chain, used for MSA generation and structural template search. The quality of the resulted structural models is characterized by the mean of per residue pLDDT (predicted Local Distance Difference Test) score (which takes values between 0 and 100, the higher value is better) and the structures are ranked accordingly^4^. The pLDDT confidence measure predicts the accuracy of the Cα Local Distance Difference Test (lDDT-Cα) for the corresponding prediction. Although this means that the high accuracy and reliability of AF2 observed in CASP14 can be transferred to predicting the structure of any protein sequences (or whole proteomes)^4,5^, this has not been validated yet and scientists do not have a clear indication how well AF2-predicted structures can be trusted. Even more, there is a special skepticism in the field of transmembrane proteins, which are challenging to investigate using either experimental or computational methods, especially because AlphaFold2 was not tuned for TM proteins. It is also not known, whether the structural model with the highest pLDDT score always corresponds to the native structure. In order to tackle these issues, we investigated whether AF2-predicted human α-helical TM protein structures exhibit correctly located TM regions. To demonstrate at a higher resolution that the predicted TM folds are native, we compared predicted structures of the ATP Binding Cassette (ABC) superfamily from the AF2-predicted 21 proteomes to existing experimental ABC folds. ABC proteins play a role in important cellular processes in all types of organisms and most of them transport substrates through the cell membrane in an ATP dependent manner^8–10^. ABCC7/CFTR is a special member, which is an ATP-gated chloride channel and includes a long intrinsically disordered regulatory R domain^11,12^. The functional form of ABC proteins is built from two highly conservative nucleotide binding domains (NBDs) and two transmembrane domains (TMDs) which can be encoded in one or separate peptide chains. The low conservation of their TMDs are related to diverse functions and their currently known TM folds are also structurally divergent and can be classified into eight groups (Pgp-, ABCG2-, MalFG-, BtuC-, EcfT-, LptFG-, MacB-, and MlaE-like folds)^13,14^. Our results demonstrate that AlphaFold2 provides reliable protein structures also for transmembrane proteins and can solve many issues associated with transmembrane protein structures.

## Results

### TM helices membrane topology assignments in AlphaFold2 structures

First, we split the human AF2 structures to soluble and transmembrane sets using the HTP (Human Transmembrane Proteome) database^15^, calculated the mean pLDDT score for each protein, and plotted their distribution (Fig. 1A and Fig. S1). Mean pLDDT values were also calculated separately for the TM and non-TM regions of transmembrane proteins. Intriguingly, soluble proteins exhibited a broader distribution and a significant area at lower pLDDT values compared to TM proteins. This was unexpected, since the majority of the AlphaFold2 learning set inherently included more soluble protein templates and the algorithm was not tuned for transmembrane proteins. However, correlation between low pLDDT values and disordered segments was observed^5^, thus our observation strongly suggested that more soluble proteins possess disordered regions than TM proteins. Interestingly, a very large portion of TM regions (53%) were predicted with high pLDDT scores (>90) (Fig. 1A) suggesting that AF2 captured the rules governing protein structures within the hydrophobic region.

**Fig. 1:**
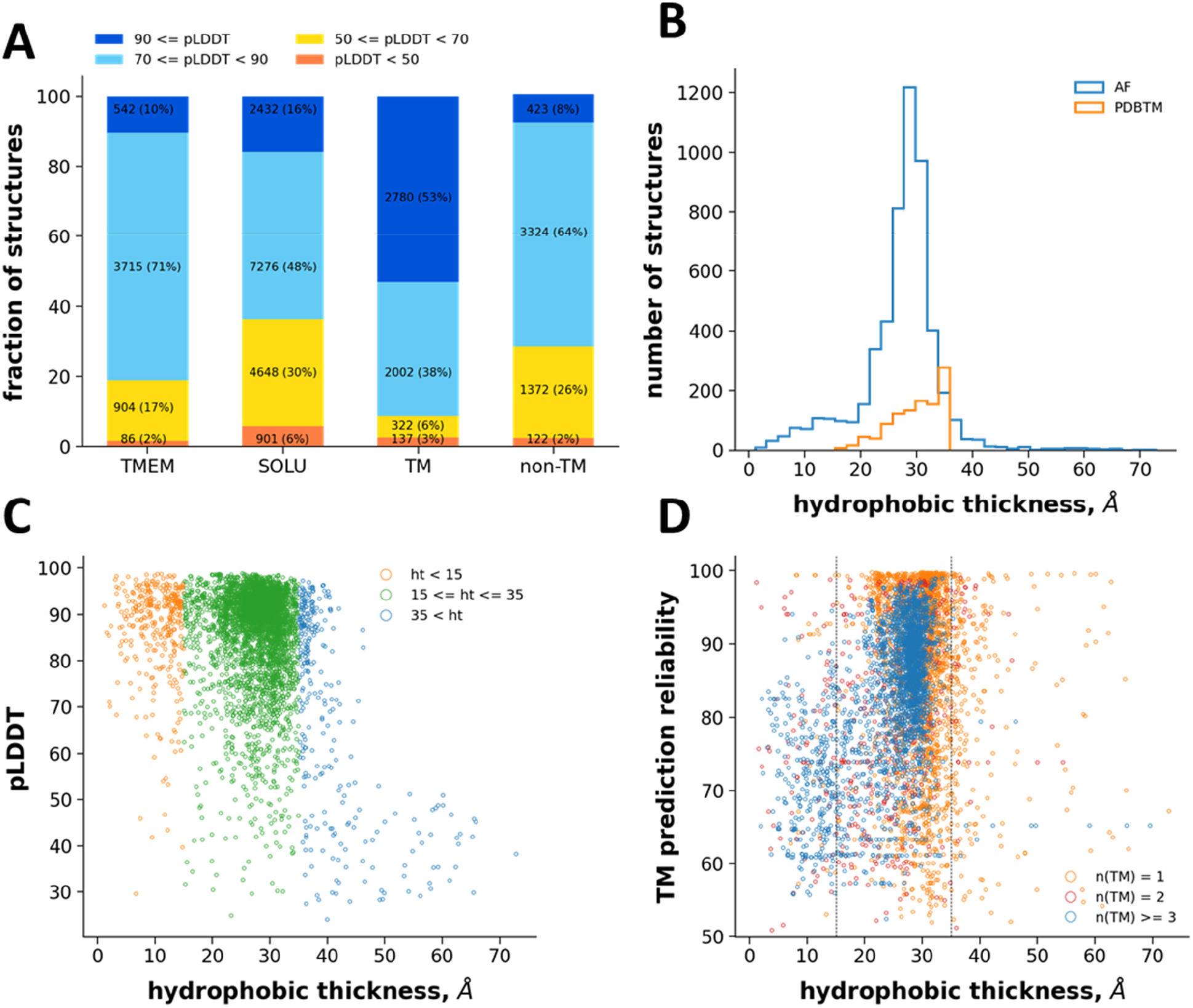
Quantitative analysis of human AF TM structures. (**A**) Mean pLDDT scores were calculated for human transmembrane (TMEM), soluble (SOLU), TM regions of TM proteins, and non-TM regions of the same proteins. The fraction of structures in reliability ranges, used in the human proteome AlphaFold2 paper^5^, are shown. (**B**) The hydrophobic thickness was calculated for human TM proteins as the distance between the center of geometry of Cα atoms of side1 and side2 of transmembrane helices. TM helices of AF2-structures were selected based on CCTOP predictions. The hydrophobic thickness of experimental structures was collected from PDBTM. (**C**) The hydrophobic thickness of each protein and the corresponding pLDDT scores were plotted. (**D**) The hydrophobic thickness of each protein and the corresponding CCTOP reliability scores are shown.

Next, we compared the spatial localization of TM helices in AF structures corresponds with rational and physiological helix orientation in a lipid bilayer slab by using the Constrained Consensus Topology prediction (CCTOP) software^16^, which includes information from both experimental and computational sources. We separated the start and end positions of predicted TM helices to two residue sets according to their localization relative to the opposite sides of the bilayer. The distance between the center of geometry of the two sets were calculated and its distribution is plotted (Fig. 1B). The majority of the membrane thickness values were in the range between 20 and 30 Å that is in the range of the hydrophobic region thickness. In order to support this finding with experimental data, the hydrophobic thickness of experimentally determined human transmembrane protein structures was retrieved from the PDBTM database^17^. The AF2 and experimental distribution largely overlapped (Fig. 1B). These observations suggested that hydrophobic thickness values below 15Å and above 35Å may indicate an erroneous AF2 structure (12%, 725 out of 5952, Table S1). We also investigated the distribution of pLDDT scores versus hydrophobic thickness (Fig. 1C). This plot indicated that AF2 structures with non-physiological thickness values can process very high pLDDT scores, consequently, these scores alone may be insufficient to select correct TM structures in blind predictions.

An inaccurate TM topology prediction of CCTOP may provide an outlier hydrophobic thickness in the case of a correct AF2-predicted structure. The CCTOP reliability versus thickness plot (Fig. 1D) indicated that the topology of most proteins, whose AF2-predicted structure exhibited hydrophobic thickness within the 15-30Å regime, was predicted with high reliability. Structures with lower hydrophobic thickness values and high CCTOP reliability were likely inaccurately predicted by AlphaFold2, while structure predictions with lower thickness and lower CCTOP scores were located in the twilight zone. Intriguingly, we observed that some of these entries may have low topology reliability because of their existence in complexes, but AF2 predicted the monomeric form correctly (Fig. S2). This suggests that AF2 may also be used to identify and aid the correction of improper membrane topology predictions.

### The experimental TM helix packing of ABC transporters overlaps with AF2-predictions

Structures of ABC superfamily members are an excellent choice to investigate AlphaFold2 performance on TM proteins, since the currently available 51 PDB entries including ABC transmembrane domains are diverse and can be classified into eight different folds represented by PDB structures (Fig. S3)^13,14^. For assessing AF2 TM protein predictions at a higher resolution, we aimed to compare AF2-built ABC TM folds with experimentally determined folds.

In order to select ABC structures from the 21 proteomes with AF2 predictions, a stringent PFAM search was performed with 29 PFAM Hidden Markov Models (Table S2) that resulted in 1,126 hits. For assessing the similarity of structures to the eight selected reference folds (Fig. S3), we employed Template Modeling score (TM-score)^18,19^. TM-scores below 0.5 indicate unrelated structures, while and above 0.5 roughly the same fold^19^. We calculated TM-scores between the 1,126 AF2-predicted transmembrane ABC structures and the eight reference structures. The best out of eight scores were saved for each structure. We found that 99.5% of the TM-score values were above 0.5 (Fig. 2A). Five out of the remaining six structures with lower TM-scores (O69723, P33359, Q2FVH1, Q2FVE9, Q2FVG9) included MalFG-like folds, but the scores were low because of their variable number of TM helices and possible disordered regions. One protein (Q2G2E2, Fig. 2B), which matched the YitT_transmembrane PFAM entry, was somewhat similar to the aquaporin/GlpF fold (e.g. PDBID: 1FX8) and suggested that the YitT_transporter PFAM entry is wrongly classified. Indeed, this fold belongs to the non-ABC, Novobiocin Exporter (NbcE) Family in the Transport Classification Database^20^. In addition to discovering potentially new folds, AlphaFold2 can aid predicting the structural class of PFAM families, such as five families out of the ABC transmembrane HMMs (Table S2).

**Fig. 2:**
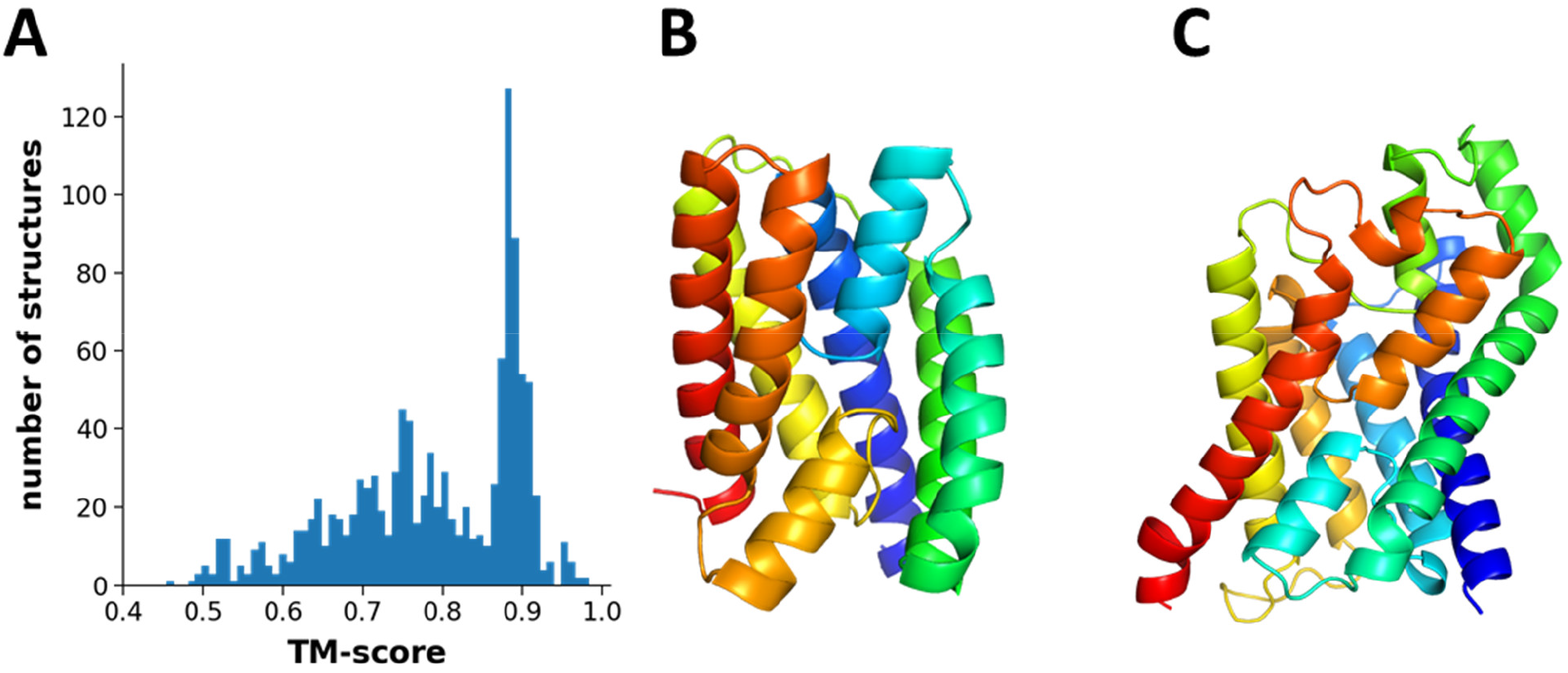
All AF2-predicted ABC structures exhibit valid ABC TM folds. (**A**) The best TM-score for every ABC TM structure from the 21 organisms calculated against every 8 ABC reference folds were selected and plotted. (**B**) The AF-Q2G2E2 fold with a TM-score lower than 0.5. This YitT fold is annotated incorrectly in PFAM and is not an ABC fold. (**C**) The Aquaporin/GlpF fold, which is somewhat similar to YitT fold and is represented by PDBID: 1FX8, possesses six TM helices and two reentrant helices forming a full TM helix-like structure.

Some of the predicted ABC structures included two additional N-terminal TM-like helices, which were somewhat distant from the core TM domain and likely are membrane associated regions, such as the L0/Lasso motif of ABCC proteins^21–23^. In many cases, membrane interacting regions, loops, and mobile regions not resolved in experimental structures have been rationally modeled by AF2, based on visual inspection (see below and Fig. S2). Thus the AF2 machine learning method clearly grasped some knowledge on a lipid bilayer around TM proteins.

### AF2 provides dimers, MD-stable structures, and hints for flaws in experimental structures

Since AlphaFold2 also used templates for structural modeling and some of the resulted structures may be considered as advanced homology models, we also performed AF2 modeling of highly studied human ABC proteins with disabled template usage.

Our targets included half transporter ABCG proteins, which consist of an NBD and a TMD in a polypeptide chain and function in homodimeric or heterodimeric complexes^14^. The first experimentally determined ABCG2-like fold was the X-ray structure of the ABCG5/ABCG8 heterodimer (PDBID: 5DO7) published in 2016^24^. Our first observation with the AF generated ABCG8 structure was regarding its soluble NBD. After the publication of the first ABCG2 structure^25^, structural alignment and sequence analysis indicated a registry shift in the first β-strand of ABCG8 NBD (Fig. 3A) that happened because of the low resolution of this region. Although AlphaFold2 exploited the 5DO7 structure as a template, the AF2-predicted ABCG8 structure did not have this error (Fig. 3A). An ABCG5/ABCG8 structure with a correct registry was also released on 07/04/2021 (PDBID: 7JR7^26^), but AF2 template search^5^ used PDB70 downloaded on 10/02/2021.

**Fig. 3:**
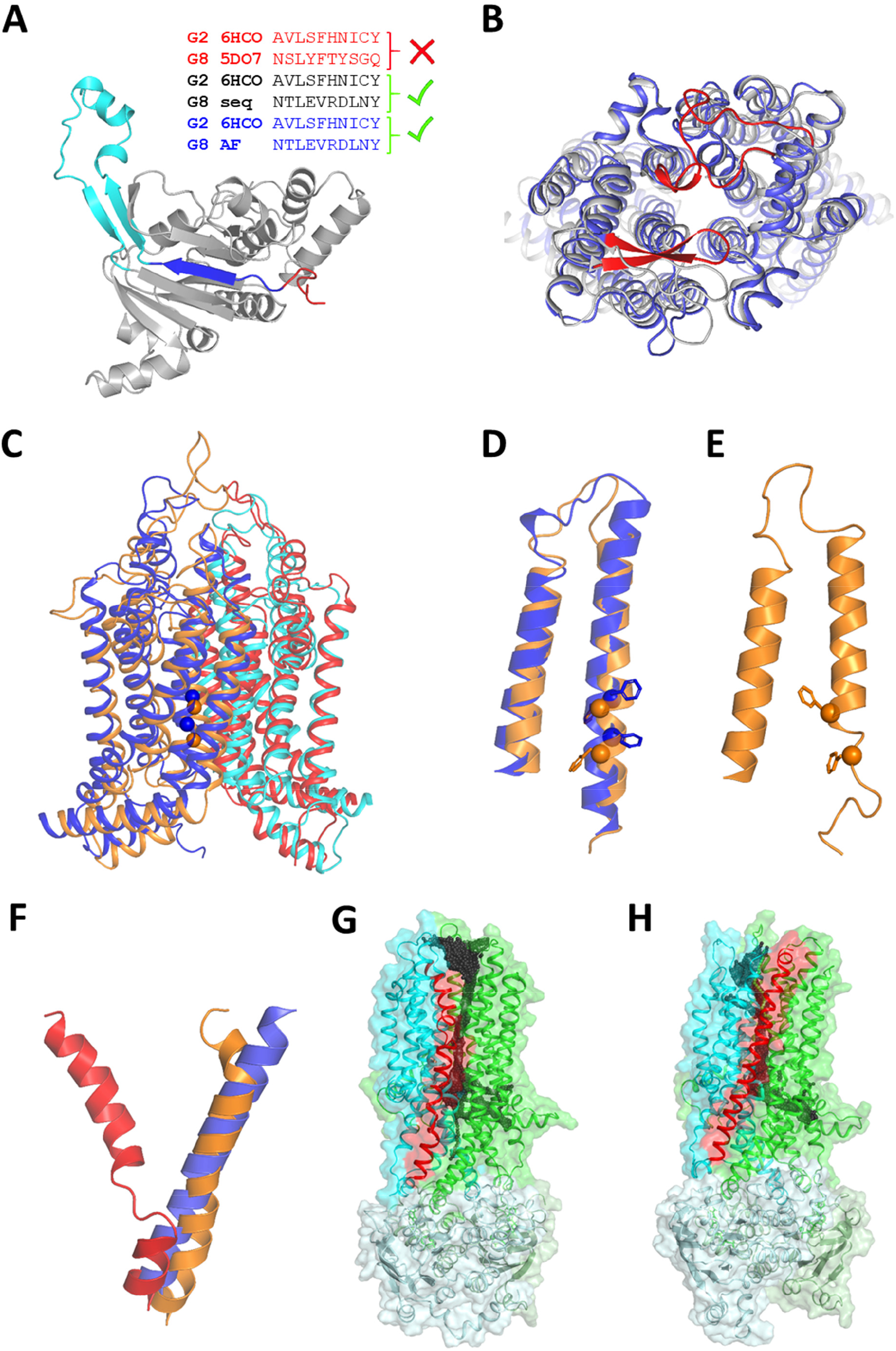
AF2 seems to handle ABC structure associated issues correctly. (**A**) ABCG2 and ABCG8 NBD β1 strand sequence alignments generated by structural alignment of 6HCO (ABCG2) and 5DO7 (ABCG5/ABCG8), by ClustalW with manual adjustment of ABCG2 and ABCG8 sequences based on ABCG2 structures, and by structural alignment of ABCG2 and AF2-predicted ABCG8 NBDs. Structure: AF2 ABCG8 NBD, blue: β1 strand, red: the segment corresponding to the β1 strand in the registry shifted 5DO7 NBD, cyan: gating loop or regulatory insertion. (**B**) Structural alignment of 5DO7 (gray) and AF2-predicted (blue) ABCG5/ABCG8 TM domains (top view). Non-conserved loops with low-quality predictions are red. (**C**) Aligned homology (orange: TMD1, red: TMD2) and AF2 (blue: TMD1, cyan: TMD2) models of AtABCG36. Blue and orange spheres label F589 and F592 in TM2 facing the substrate binding pocket. (**D**) The magnified view of AtABCG36 TM1 and TM2 indicates that the alignments are not shifted but that spatial localization and side chain packing differ. (**E**) TM2 in the homology model unwinds in MD simulations. (**F**) zfCFTR TM8 is kinked in PDBID:5W81 (red) along with other structures and it is straight in both MRP1-based model (orange) and AF2-predicted structure (blue). The helices are extracted for visualization from a full TM domain alignment. (**G**) Surface representation of zfCFTR (PDBID:5W81). Red: TM8, green: TMD1, cyan: TMD2, pale green: NBD1, pale cyan: NBD2, black spheres: CAVER spheres indicating channel opening towards the extracellular space and the extracellular boundary of the lipid bilayer. (**H**) Surface representation of zfCFTR with MRP1-modelled, straight TM8. No lateral opening to the extracellular membrane leaflet can be observed.

To assess ABCG5/ABCG8 TMD predictions, we ran AF2 without any application of templates. First, the ABCG5 TMD predictions were of exceptionally good quality regarding the RMSD (root mean square deviation) and TM-score values of 0.78Å and 0.94, respectively, when compared to the ABCG5 chain in the 5DO7 structure. Second, we investigated ABCG5/ABCG8 heterodimer predictions. Since only single chains can be submitted to AlphaFold2, we concatenated the two sequences with a part of the CFTR R domain sequence (a.a. 675-800). This disordered sequence was sufficiently long not to constrain the conformational space of the dimer and did not exhibit strong intramolecular interactions even in its native, AF2-predicted structural environment (Fig. S4). The predicted TMD dimer exhibited 0.001Å RMSD and 1.00 TM-score value when compared with the 5DO7 structure (Fig. 3B).

To investigate if AlphaFold2 can distinguish between intra- and intermolecular interactions in the case of homomeric complexes, we performed a prediction with ABCG2, which forms homodimers^27^. The complex of the two identical TMDs was also predicted exceptionally well (2.42Å RMSD and 0.9 TM-score when compared to PDBID: 6VXF). Interestingly, cysteine residues forming intra- and intermolecular disulfide bonds were close to each other (Fig. S5).

We also examined how AF2 structural models can supplement or replace homology models in molecular dynamics (MD) simulations. The TM regions of distant ABC proteins exhibit low sequence conservation with good accordance of their dissimilar functions and substrates. However, their folds in a family are highly conserved, thus homology modeling can provide high quality homology models^28–31^. We chose AtABCG36/PEN3/PDR8^32^ from the model plant *Arabidopsis thaliana*, which is a well-investigated full transporter of the ABCG subclass for that no structures yet exist. When the homology model exhibiting two ABCG2-like TMDs (Fig. 3C) was inserted into a membrane bilayer and subjected to MD, one portion of an α-helix, which is part of the central drug binding pocket, exhibited fast unfolding in an equilibrium MD simulation. Then the AF2-predicted AtABCG36 structure under the same conditions remained stable in an MD simulation (Fig. S6).

The CFTR/ABCC7 chloride channel is also a member of the ABC superfamily with a Pgp-like fold. The functional mechanism of this protein is of interest, since some mutations effect channel gating and cause cystic fibrosis^33^. One of its structures was determined using cryo-EM under activating condition, in the presence of ATP and phosphorylation, but the extracellular pore of the channel remained in a closed state, most likely due to a kink in TM8, corresponding to an unwound segment in the transmembrane region^34^ (Fig. 3F). This kink is present in most CFTR structures (PDBIDs: 5UAK, 6O2P, 6MSM, 6O1V, 5UAR, 5W81)^34–37^. However, the kink is absent from the chicken CFTR structure (PDBIDs: 6D3S and 6D3R)^38^ and such a conformation has not been detected in other ABC structures. We performed equilibrium simulations with the 5W81 structure^12^ to detect channel opening, but appearance of tunnels with sufficient diameter to pass chloride ions were rare events and was observed only once out of 22 simulations (6×100ns + 16×35ns, 427/116,000 frames, 0.004%). Intriguingly, many of the conformations provided a tunnel opened towards lipid molecules of the extracellular membrane leaflet (Fig. 3G). After correcting the kink by homology modelling based on the MRP1 structure (PDBID: 5UJ9) (Fig. 3F), opening of the extracellular pore could be observed in 5 out of 6 simulations at a higher probability (6×100nx, 2,245/60,000 frames, 3.74%). Remarkably, modeling CFTR TMDs using AlphaFold2 without CFTR or any templates resulted in a conformation similar to that of MRP1 with a straight TM8 helix (Fig. 3F,H). Since TM8 has been suggested to be flexible regarding to its membrane embedment^39^, it is likely sensitive to its environment and based on the functional assays and the structure determination protocol^35^, the detergent added in the last step (3 mM fluorinated Fos-Choline-8) likely biased the structure.

### Prediction of a new TM fold, never seen by AlphaFold2

The most obvious test for assessing AlphaFold2’s ability to predict membrane protein structures, was running a blind prediction. For this objective we used the multiple peptide resistance factor (MprF) transmembrane domain sequence, whose structures was published this year by Song *et al*.^40^ (PDBIDs: 6LVF and 7DUW) Thus their novel TM fold (Fig. 4A) was not present in the AlphaFold2’s training set. We disabled template usage in AlphaFold2 run, since the 6LVF structure is already in the pdb70 dataset used by AF2. Since the top ranked AF2 structure was not a matching AF2-prediction (Fig. 4B), we performed the prediction several times (n=6) and compared the predicted structures to the transmembrane domain of 7DUW using TM-score. Plotting the pLDDT scores versus TM-scores (Fig. 4C) indicated that among the 25 predicted structures the one with the best pLDDT score was the most similar to the target structure (highest TM-score in the set) (Fig. 4D). In addition, this plot also included two important hints regarding evaluating predicted structures. First, AF2 structures with high pLDDT values can be highly distinct from the native fold. Second, AF2 neural network model 5 did not perform well on this membrane protein, while model 2 and model 4 even in the middle pLDDT range provided the same fold as the native. We were unsuccessful in predicting the dimeric form of MprF that was likely due to the very small protein-protein interaction interface and the fact that lipid molecules are also play a role in dimer stabilization^40^.

**Fig. 4:**
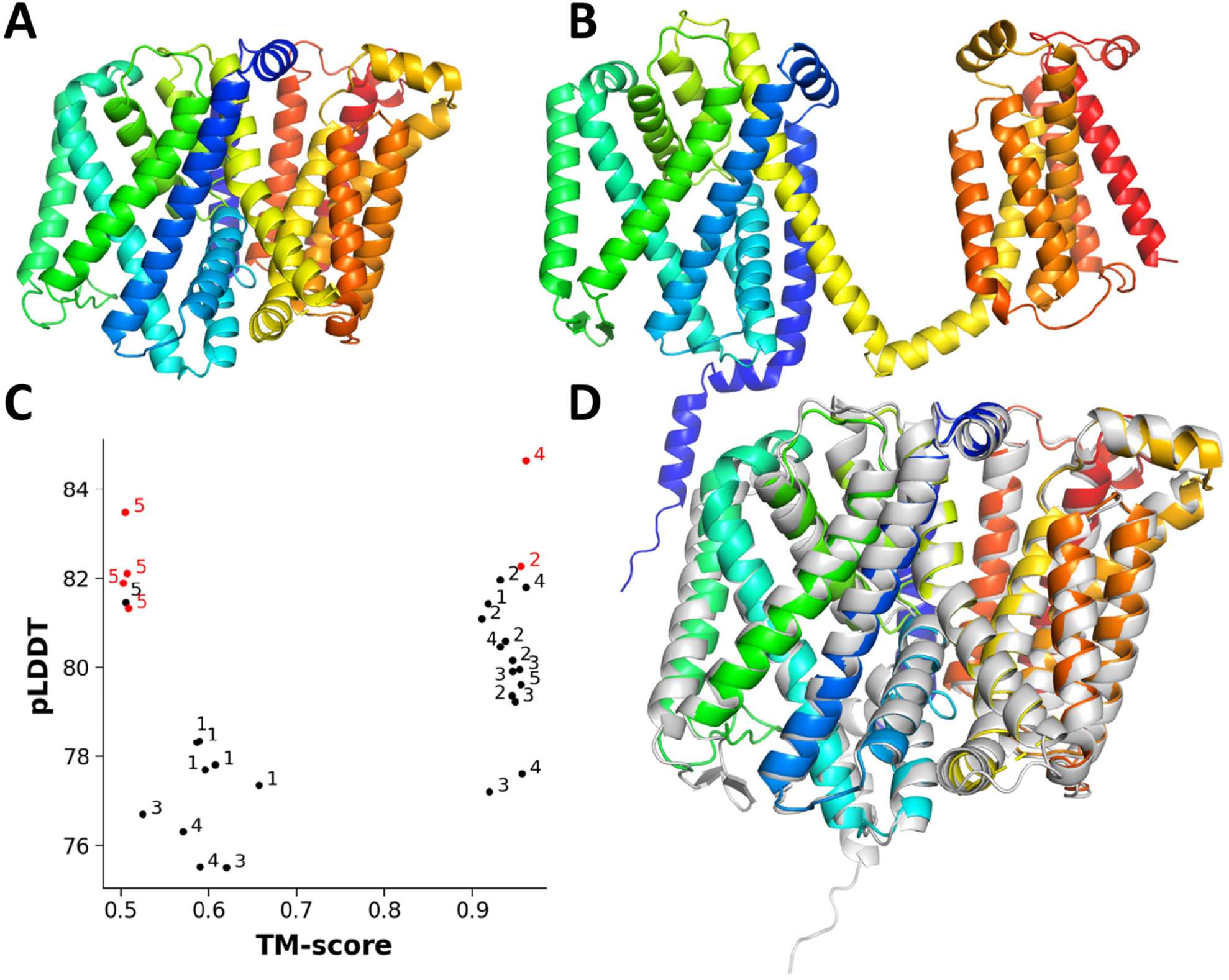
Predicting new TM folds. (**A**) The novel TM fold of MprF (PDBID: 7DUW). Blue to red: N- to C-termini. (**B**) An AF2-predicted fold that significantly differs from the experimentally determined fold (e.g. N-terminus starts on the opposite side of the membrane). (**C**) pLDDT and TM-score values, calculated for every structural model from six runs, were plotted. Numbers (1-5) indicate the corresponding AF2 models. Red points were the top ranked hits from a given run. (**D**) Structural alignment of the experimental structure (rainbow colors) and the prediction with the best pLDDT score (gray).

## Discussion

We demonstrated that at least ∼90% of the AF2-predicted TM structures of the human proteome represented membrane-protein like structures, using the most available and reliable measure, the location of TM helices from consensus predictions and experimental structures, for assessing TM protein structure quality at a large scale. While the pLDDT score distribution did not shift much to lower values compared to soluble proteins (Fig. S1) it is likely valid to state that AF2 predict TM proteins as good as soluble proteins. However, predicted TM structures with low hydrophobic thickness and high pLDDT score (Fig. 1C) and our predictions with the novel MprF fold (Fig. 4C) suggest that evaluation depending solely on pLDDT score may not be sufficient to select the best AF2-predicted model, at least in the case of TM proteins. A similar conclusion was drawn comparing the AF2-predicted and cryo-EM structures of a pump-like channelrhodopsin with structural features never seen before^41^. Based on our results, the quality of AF2 models can also be investigated by MD simulations in low-throughput studies, since low-quality AF2 models should be as instable as incorrect homology models (Fig. S6) or experimental structures^42^.

Our results demonstrate that AlphaFold2 is a highly valuable tool in many areas of TM protein research. It can highlight proteins with potentially new folds (Fig. 2B) and select them for experimental structure determination, resulting an increase in the experimental fold space. AlphaFold2 can also associate PFAM families with structural folds that will aid functional annotation of yet uncharacterized proteins (Table S2). Importantly, the AF2-predicted AtABCG36 structure revealed that AF2 in combination with a structure alignment method (e.g. TMalign) can support any methods using sequence alignments. The stability of TM2 of AF2 AtABCG36 model in MD simulations strongly suggests that the AlphaFold2s model building and relaxation protocols provide valid inputs also for drug/protein interactions studies.

AlphaFold2 is being suggested to be exploited in molecular replacement protocol aiding experimental structure determination^43^. One of our results, that the register shift in ABCG8 NBD is corrected by AF2 (Fig. 3A), supports this type of applications, and also suggests that comparison of PDB structures with the corresponding AF2 structures may detect structural errors and contribute to improvement of PDB quality. This observation of corrected registry shift in the presence of an incorrect template and the absence of the kink in CFTR TM8 upon disabling template usage, are strong indications that the neural network behind AlphaFold2 is not overfitted and it can overcome memory footprints originating from training.

We also demonstrated that AF was capable to predict transmembrane dimer structures independently of their homo- or heteromeric nature (Fig. 3B and Fig. S5). Though, this success may be at least partially resulted by the footprint of these complexes themselves in the AF2 neural network. Interestingly, AF2 was not trained for multimer predictions and it was also reported being successful for protein-peptide docking^44^, in which peptides were not involved in alignments. These observations suggest that AF2 “learned something more” than protein structure and something deeper on the mechanism of protein folding. Since knowledge on the folding steps and intermediate structures is important to understand and cure folding diseases^45,46^, deciphering hidden information on folding directly or indirectly from AF2 neural network is an important future objective.

In summary, our study underscores that AlphaFold2 provides reliable protein structures also for transmembrane proteins and we demonstrated its unexpected performance in many areas associated with transmembrane protein structures. The artificial intelligence inside AlphaFold2 can predict various structural information and correct structure related flaws (e.g. registry shift, alignments, TM topology prediction, etc.) same as or better than humans.

## Methods

### Databases and associated software

AlphaFold2 structures predicted for 21 proteomes were downloaded from https://alphafold.ebi.ac.uk in July, 2021. Proteins and their structures are identified in the manuscript with their UniProt accession number. Human Transmembrane Protein database^15^ (02.06.2021) was received as an XML file from http://htp.enzim.hu. The data also contained CCTOP^16^ (http://cctop.enzim.ttk.mta.hu) predictions and their reliability values. The hydrophobic thickness of experimentally determined human TM protein structures was retrieved from the PDBTM database (http://pdbtm.enzim.hu,2021-07-23)^17^. Python was used to parse their XML files.

ABC PFAM entries were identified at https://pfam.xfam.org (n=29) and extracted from the Pfam-A.hmm file. The selected entries and their accession numbers are listed in Table S2. The sequence of every AF2 structure was searched using HMMER hmmsearch (http://hmmer.org)^47^. The E parameter was set to 0.001 and the match length was restricted to a minimum of 90% of the HMM profile length. The hmmsearch output was parsed using BioPython^48^.

### Data analysis and visualization

MDAnalysis^49^ and NumPy^50^ Python packages were used for calculation of mean pLDDT values and hydrophobic membrane thickness. The pLDDT value of each residue were extracted from the B-factor column of AF2 structure files. For TM thickness calculation end positions of TM helices were retrieved from HTP/CCTOP and divided into two groups representing the two sides of the membrane. Plotting was done with Matplotlib (https://matplotlib.org)^51^.

TM-score was calculated with TMalign^52^. Reference ABC structures are listed and shown in Fig. S3. Their TM domains were selected manually.

Molecular visualization and RMSD calculation were performed using PyMOL (The PyMOL Molecular Graphics System, Version 2.4.0 Schrödinger, LLC). RMSD of MD trajectories was calculated with the GROMACS rms tool.

### Running AlphaFold2

AlphaFold2 was downloaded from github and installed as described (https://github.com/deepmind/alphafold) on a Debian 10 box with an AMD Ryzen Threadripper 2950X 16-Core Processor. 96GB RAM was installed and ∼75GB peak usage was observed during jackhmmer run. The calculation was accelerated by an NVidia Quadro P6000 GPU with 24GB RAM, which was almost fully utilized when the predicted sequence length was 1,571. The required databases were located on two 2TB HDD in a RAID0 setup. Typical run timings were: “features”: 25-60 min, “predict_and_compile_model_*”: 3-50 min, “relax_model_*”: 1 min - 6 h based on input sequences between 290 and 1,571 a.a. length.

In order to exclude CFTR structures as templates from predictions, we modified run_alphafold.py, docker/run_docker.py, and alphafold/data/templates.py scripts to implement a -skip function. The modified scripts can be downloaded from http://alphafold.hegelab.org. Template usage was disabled by setting --max_template_date option to 1900-01-01. Dimer predictions were run by concatenating sequences with a part of the intrinsically disordered CFTR R domain, a.a. 675-800. pLDDT scores and ranking of predicted structures were extracted from the ranking_debug.json file.

### Homology modelling

AtABCG36 (UniProt ACC: Q9XIE2) was homology modeled based on an ABCG2 homodimer structure (PDBID: 6HZM) using Modeller^53^. Sequence alignment was generated using ClustalW^54^ and adjusted manually. One hundred structures were generated and the one with the best DOPE score was selected for MD simulations.

zfCFTR TM7 and TM8 was homology modeled similarly. The two helices were set for modelling based on the corresponding regions of MRP1 (PDBID: 5UJ9^23^) and the rest was kept static and based on the 5W81 zfCFTR structure.

### Molecular dynamics simulations

MD simulations with AtABCG36 were performed using GROMACS 2019 with the CHARMM36m force field^55,56^. Simulation systems were prepared using CHARMM-GUI^57,58^. Structural models were oriented according to the OPM (Orientations of Proteins in Membranes) database^59^ and all N- and C-termini were patched with ACE (acetyl) and CT3 (N-Methylamide) groups, respectively. The proteins were inserted in a bilayer with 1:1 POPC:PLPC (1-palmitoyl-2-oleoyl-sn-glycero-3-phosphocholine: 1-palmitoyl-2-linoleoyl-sn-glycero-3-phosphocholine) in the extracellular leaflet and 45:40:10:5 POPC:PLPC:POPS:PIP2 (POPS: 1-palmitoyl-2-oleoyl-sn-glycero-3-phospho-L-serine, PIP2: phosphatidylinositol 4,5-bisphosphate) in the intracellular leaflet. Both systems with the homology model or the AF2 structure were energy minimized using the steepest descent integrator (values for max. steps 50,000 and max. force 500 kJ/mol/nm were set). Six equilibration steps, according to the standard CHARMM-GUI protocol, were applied with decreasing position restraints. In the production run, Nosé-Hoover thermostat and Parrinello-Rahman barostat with semiisotropic coupling were employed. Time constants for the thermostat and the barostat were set to 1 picosecond and 5 picosecond, respectively. The fast smooth PME algorithm^60^ and LINCS algorithm^61^ were used to calculate electrostatic interactions and to constrain bonds, respectively. GROMACS rmsf tools were used to calculate RMSF (root mean square fluctuation).

Simulations with the zfCFTR structure containing the kinked TM8 have been published and the protocol and parameters were described there^12^. The structure with the straightened, MRP1-based TM8 was subjected to MD simulations using the same protocol, including the same version of GROMACS, force field, and lipid composition. Channel pathways were determined using CAVER^62^ as described^12^.

## Data Availability

All input data are available from public resources and their accession numbers are listed.

## Code availability

Modified AlphaFold2 scripts can be downloaded from http://alphafold.hegelab.org.

## Acknowledgements

We thank H. Tordai, R. Padányi (Semmelweis University, Hungary) and G. Gyimesi (University of Bern, Switzerland) for their helpful suggestions. We acknowledge the computational resources made available B. Babics (Boblem IT Co.), Governmental Information-Technology Development Agency (https://hpc.kifu.hu), the Grubmüller laboratory at Max Planck Institute (https://www.mpibpc.mpg.de/grubmueller), and Wigner GPU Laboratory (http://gpu.wigner.mta.hu) and we thank the help of their members. This work was supported by funds to T. Hegedűs from the Cystic Fibrosis Foundation (CFF): HEGEDU20I0 and from NRDIO/NKFIH: K127961; to G. Lukacs from CCF LUKACS20G0, CIHR, CFI and Canada Research Chair Program to G. Lukacs; to M. Geisler from the Swiss National Funds (310030_197563).

## Author contributions

TH, MG, and GL conceived ideas and wrote the manuscript. TH and BF performed calculations and their analysis.

## Materials & Correspondence

should be addressed to TH.

