## Supplementary Material for "AlphaFold2 transmembrane protein structure prediction shines"

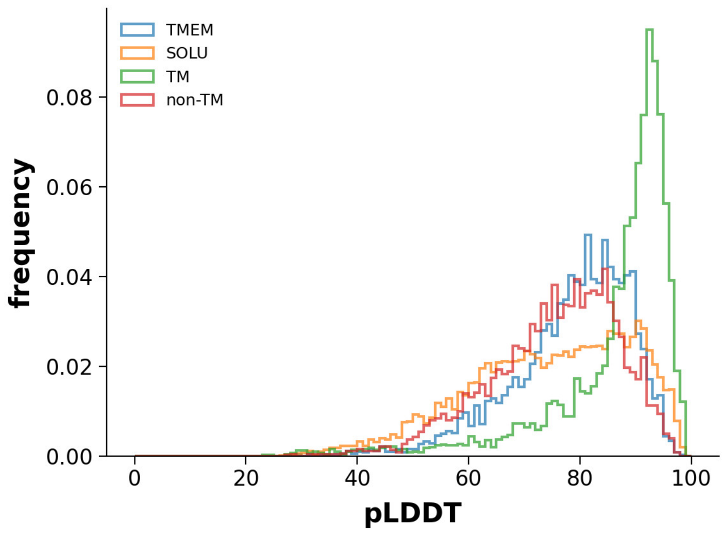


**Fig. S1: Mean pLDDT distribution in various subsets of human AF2-2 predicted protein structures.** TMEM: transmembrane proteins, SOLU: soluble proteins, TM: transmembrane region of TM proteins, non-TM: non-transmembrane regions of TM proteins.

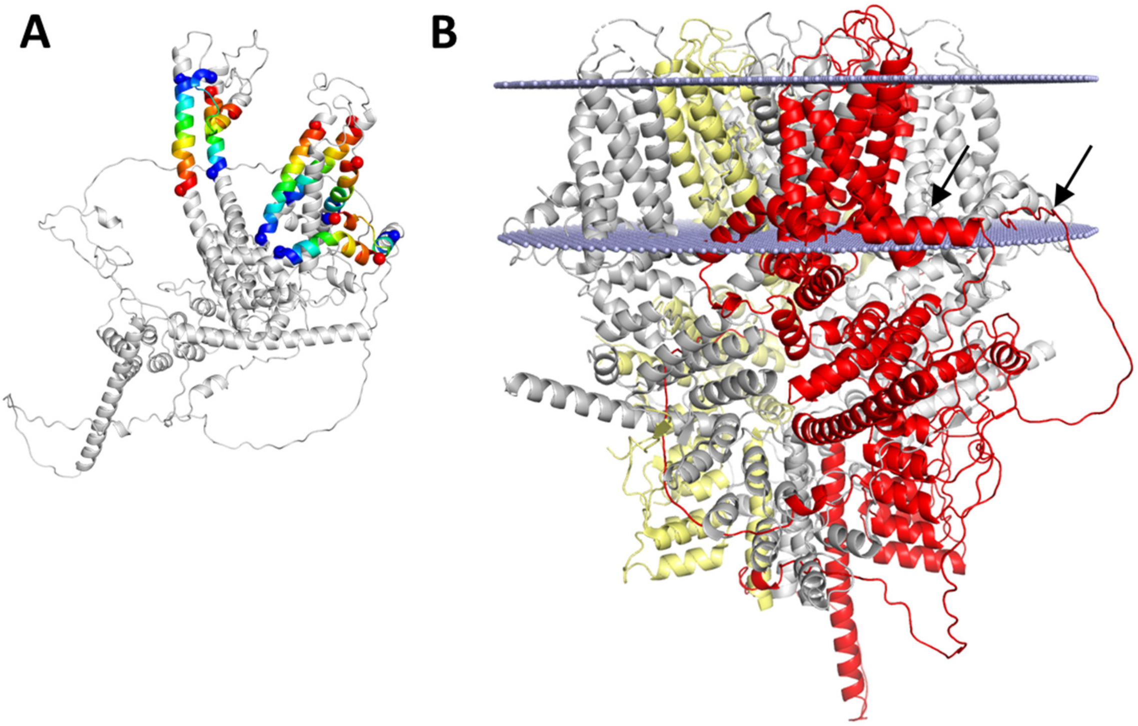


**Fig. S2: Low thickness and low CCTOP reliability values may indicate structures forming complexes.** (**A**) The TRPC4 (short transient receptor potential channel 4, ACC: Q9UBN4) exhibits low thickness value and some of its TM helices in the AF2 structure do not pass the membrane, but one forms a reentrant helix and one another is located parallel to the bilayer plane, based on CCTOP prediction with a reliability score of 63. Transmembrane helices based on CCTOP prediction are colored from blue to red from side1 to side2. End positions are labeled with spheres. (**B**) The structure of mouse Trpc4 (PDBID: 5Z96) and its chain A is replaced with human AF2 TRPC4 structure (red, RMSD: 0.6Å). Light gray: chain B, yellow: chain C, gray: chain D, light blue: membrane bilayer predicted by OPM (https://opm.phar.umich.edu). Arrows indicated regions, which are not present in the experimental structure and do not immerse irrationally deep into the membrane region. Thus AlphaFold2 may have some idea about the presence of membranes around TM proteins.

| fold class | reference PDB |
|---|---|
| Pgp-like | 4M1M |
| ABCG2-like | 6HCO |
| MalFG-like | 2R6G |
| BtuC-like | 4FI3 |
| EcfT-like | 4HUQ |
| LptFG-like | 5X5Y |
| MacB-like | 5LJ7 |
| MlaE-like | 7CH0 |

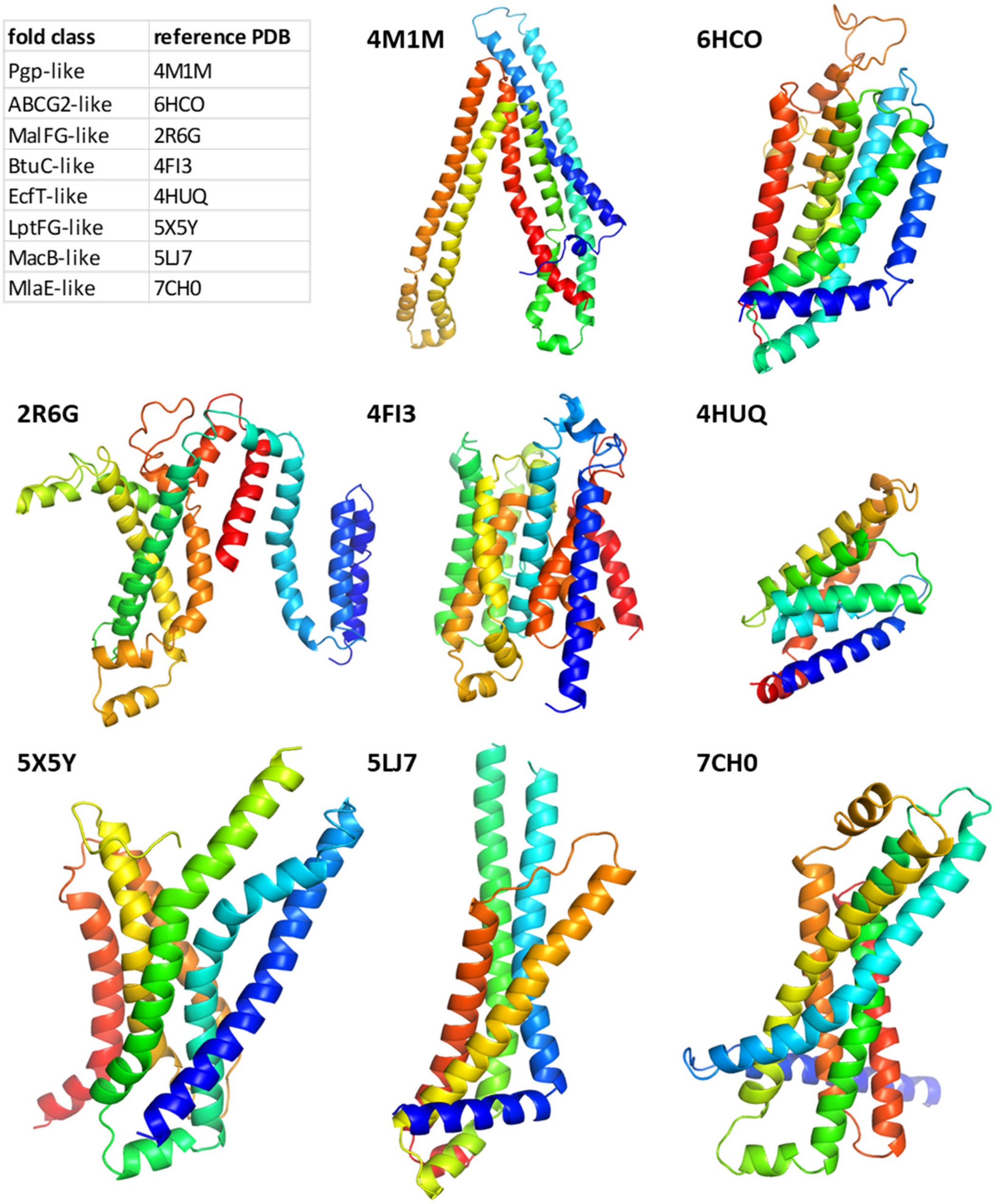


**Fig. S3: Structures used as reference ABC transmembrane domain folds.** One TMD structure was extracted from a structure from each fold class. These TMD structures were used as reference folds for classifying TMD domains using TMalign.

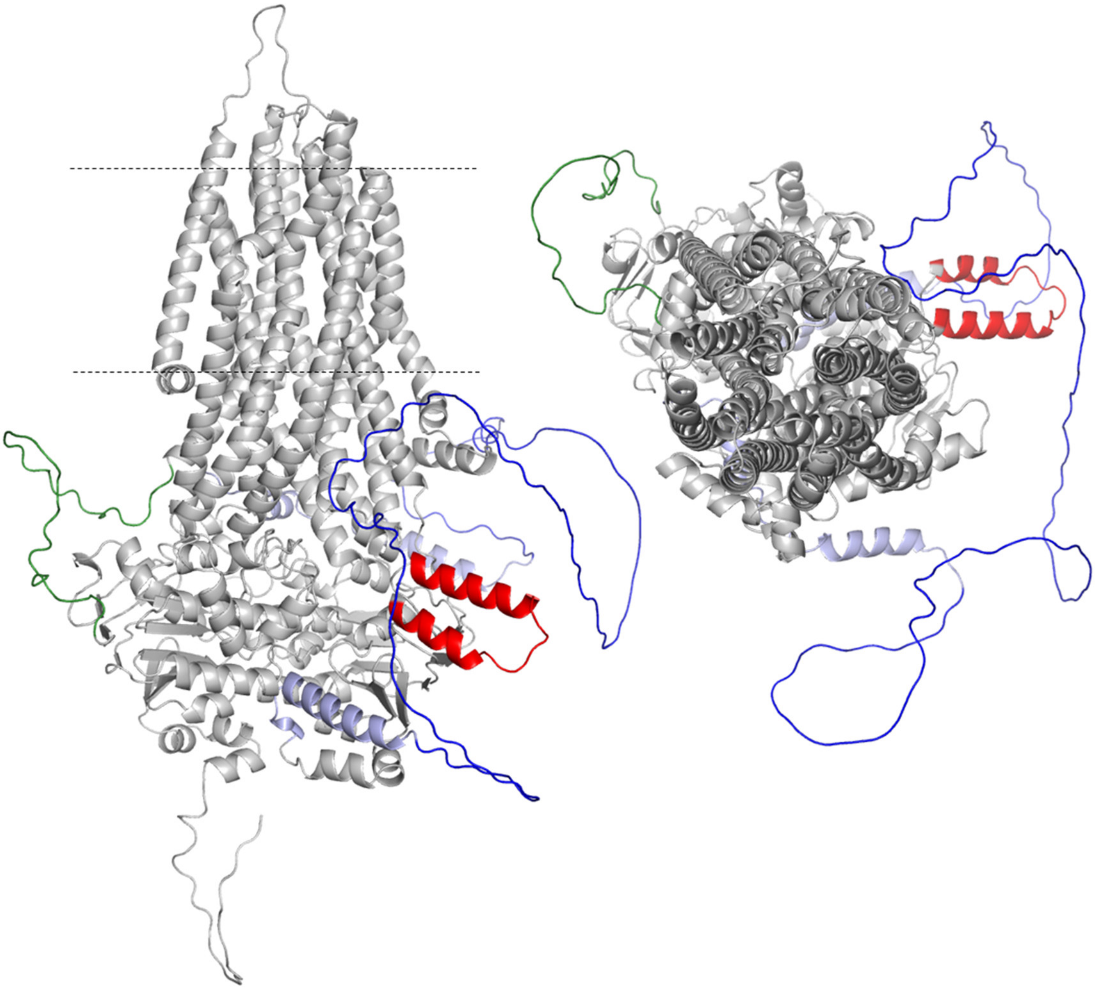

**Fig. S4: The AF2 structure of human CFTR (ACC: P13569).** Side view and top view from the extracellular space. Blue and green: disordered regulatory domain and linker region, respectively; none of them enters the hydrophobic bilayer region which is indicated by dashed lines; red: dynamic regulatory insertion, which is located in the first nucleotide binding domain and is only partially resolved in isolated NBD1 or full-length structures.

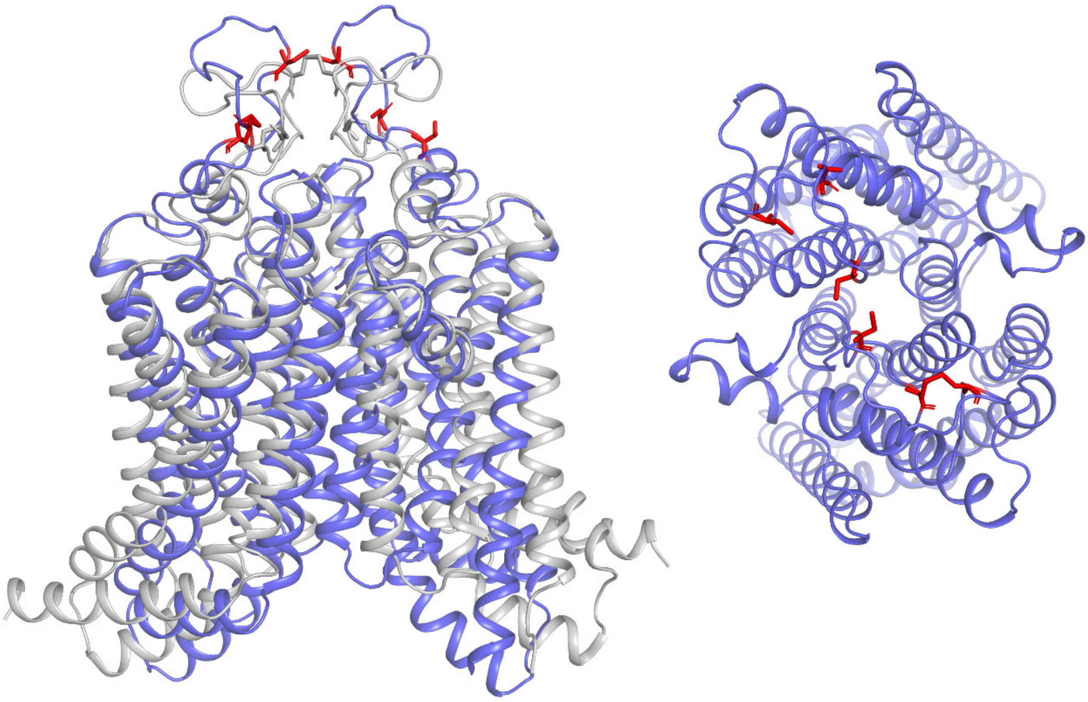

**Fig. S5: ABCG2 homodimer.** Two ABCG2 (Q9UNQ0) TMD sequences were linked with a CFTR R domain segment and subjected to AlphaFold2 prediction without template usage. Side and top views from the extracellular space, the R domain segment is not shown. Grey: experimental structure (PDBID: 6HCO), blue: AF2 model, red sticks: AF2 cysteine residues forming intramolecular and intermolecular disulfide bonds, located in the extracellular loop 3.

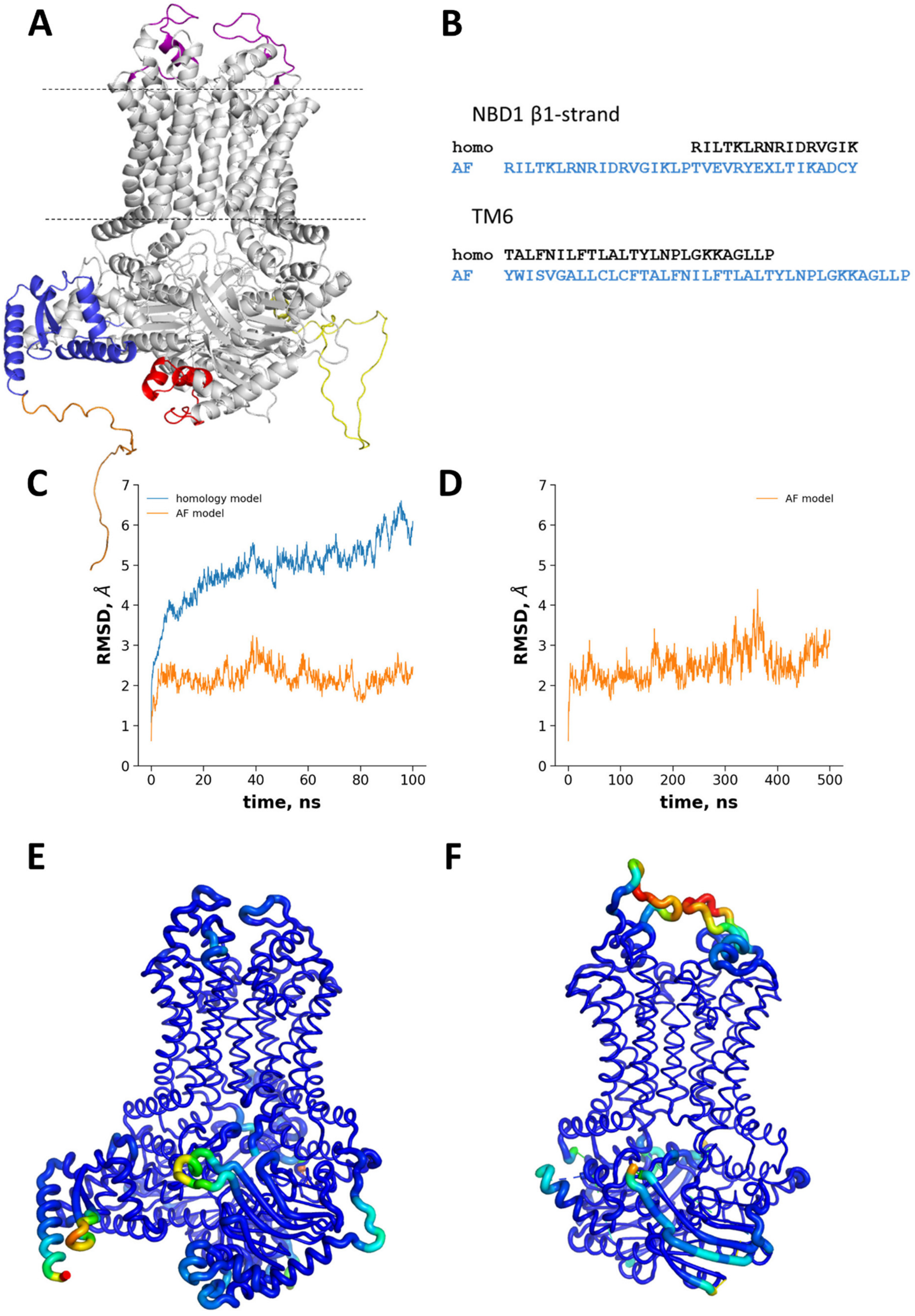
A
B
NBD1 β1-strand
homo RILTKLRNRIDRVGIK
AF RILTKLRNRIDRVGIKLPTVEVRYEXLTIKADCY
TM6
homo TALFNILFTLALTYLNPLGKKAGLLP
AF YWISVGALLCLCFTALFNILFTLALTYLNPLGKKAGLLP
C
homology model
AF model
RMSD, Å
time, ns
D
AF model
RMSD, Å
time, ns
E
F

**Fig. S6: Comparing MD simulations with the homology model and the AtABCG36 AF2 model (ACC: Q9XIE2).** (**A**) AF2 provided structures for regions that were difficult (non-conserved extracellular loops #3 and helices between NBD and the linker, labelled by purple and red, respectively) or impossible (N-terminal helices, blue) to homology model. (**B**) We misaligned the non-conserved β1 strand in the homology model, which segment is also registry shifted in ABCG8 experimental structure (Fig. 3A). TM6 was misaligned because the proceeding non-conserved EL3. (**C**) The homology model highly deviated from the initial structure more than the AF2 model based on RMSD (root mean square deviation). (**D**) The AF2 structure exhibited stability also on longer time scales (500 ns). (**E**) The homology model exhibited fluctuations in EL3 and β-strands of NBD1 higher than observed for the AF2 model in MD simulations (**F**). Warmer color and larger thickness indicate higher B-factor, which was calculated by GROMACS rmsf tool for frames between 50 and 100 ns. The disordered regions a.a. 1-39 and 809-863 of AtABCG36, were removed before MD system building, since they exhibited lower quality score and could have possibly interfere with the simulation.

**Table S1: Potentially erroneous AF-predicted human TM structures with hydrophobic thickness lower than 15Å or higher than 35 Å.**

| | UniProt ACC |
|---|---|
| **d < 15 Å**<br>416 models<br>350 proteins | Q01668, Q9NSA2, Q7Z4N2, Q9Y2P4, Q7Z7M0, O43462, Q8IWY9, Q9NUH8, Q8IZQ1, Q2QL34, P21817, Q5BJD5, Q9NZV8, Q8WW34, Q96N23, Q04671, Q96T54, Q13563, Q9H6D3, Q3KNW5, Q86UB9, Q9H5I5, Q5K4L6, Q14534, Q9UKG4, Q9Y4D8, Q92521, O75298, Q8IVB4, Q13507, Q5BKT4, Q13571, A5PLL7, Q9UL62, P31513, A8MWL7, O14735, Q9BX84, P48651, O95180, O14880, Q9Y2E8, Q96P56, Q7RTX7, Q5JUK3, A6NKG5, Q92819, Q15858, Q8NBR0, Q9NUD9, Q9UBN4, Q9NV58, Q9BTD3, Q9P055, A5PLK6, Q15878, Q92575, Q9BQB6, Q9Y2U2, Q6PP77, Q92952, Q96BM0, Q14573, Q9Y6M7, P51811, P51795, O95406, Q75T13, P22459, Q96L42, Q6IF36, Q9BXS9, Q8WUD6, P42858, P36269, P40879, Q9P2D8, A8MWK0, P38435, Q53FP2, Q2VPB7, Q9UK17, Q9Y4A5, Q9BVK8, Q96F25, Q99466, Q6UX98, O14524, Q6UVM3, Q14689, O95279, Q6P1A2, Q7Z2W7, Q5GH77, Q12908, Q5W0B7, Q9Y210, Q14674, Q9H0A0, Q9NX78, O00555, B2RXF0, Q9Y5Y9, O94919, A1A4F0, Q86VD9, Q6NUI2, Q6ZRR5, Q7Z5M5, Q9UI33, Q9UBU6, Q8NGG1, O15056, Q6ZWT7, Q8TEB9, Q6ZN68, P30988, Q9Y2K9, Q5I7T1, Q4G0N8, Q9BY78, Q9H2B4, P48995, Q9BWW8, Q5VTY9, Q9NRC1, Q96GC9, Q7Z449, Q6NTF9, Q8TEQ8, Q9H3K2, P46977, Q92604, Q2LD37, Q9NXE4, Q9UBY0, Q96JW4, Q16799, Q96N66, Q96Q80, Q92543, Q9UQ05, Q9NVV0, Q00975, Q96PR1, Q9Y672, Q5TEA6, O43511, A6NM10, P35499, P0C7U9, Q8IZY2, Q6PI25, P51787, Q13488, Q9NRR6, Q6ZUK4, Q9BVC6, C9JQI7, Q8NCM2, Q6NXT6, Q8IVJ1, Q14667, Q9P0X4, Q8NFU1, Q6PCB7, P48751, Q6ZXV5, Q8WWI5, O95197, Q9UI14, O00219, O43525, Q13936, Q9HB14, A6NKP2, Q6P4A7, Q8N1M1, Q86WA9, Q96B96, Q7Z3C6, Q9NR77, O76024, Q7LBE3, Q9Y257, Q6ZNC8, Q9UH99, Q9NR82, P50443, Q6TCH4, Q71RG4, Q8IU89, Q9UGP8, Q96HV5, O60840, P61550, Q01118, P39210, Q8IY26, A0A1B0GVZ9, Q9UKU0, Q92839, O43526, Q5T197, Q12791, Q9H0H0, H0YL14, O94911, Q5VW36, Q6P1M0, A6NFY4, P31213, Q9P003, Q8N5T2, Q7Z5H4, Q8NBV4, Q9BVG9, Q96NB2, Q86UL3, Q9P0S9, P35498, Q9UMS5, Q8N4W6, Q96G23, Q7Z418, Q9H330, Q7L1W4, Q9BV10, Q9NY46, O60427, Q6ZVX9, Q9H6F2, Q99250, Q6ZMZ0, Q8TDI7, Q13224, Q14656, Q9BX79, Q03721, Q86UQ4, P35610, Q8TBE1, O14649, Q13535, Q8IUA7, Q9ULQ1, Q01740, Q14524, Q9H490, Q9H427, Q14003, Q9BRB3, Q9H9B4, P58743, Q2T9J0, Q9UQD0, Q8TCJ2, P40305, Q86X29, Q6P7N7, Q10981, Q9NT68, Q9UG22, Q58HT5, Q6N022, P59826, Q6UWV2, Q5JXX7, Q99942, Q6Y2X3, H3BN30, Q96SE0, Q68CQ1, Q15392, O15013, Q86SJ2, Q7Z407, Q96C03, B6SEH9, Q16836, Q5TGY1, O75445, Q96AD5, Q9UKZ4, Q86XA9, Q6ZV70, Q9UJ14, Q9ULC5, P60509, Q8IXK2, Q68CK6, P0DMW4, Q9H1A4, Q9NSK7, Q8WXF7, Q9P265, Q8TCG1, Q2TAA5, Q96GF1, P54829, O43914, Q8IYT2, P09110, Q99732, Q9H267, Q3SXM5, Q08AH3, Q9NUB1, A6NEH6, P51841, O75871, Q9P0S3, Q902F9, Q86V21, Q8N414, Q6GMR7, Q86VF5, O95237, Q6GPH6, O95976, Q6P1Q0, Q96MZ0, Q9NZC3, P61565, Q8N9F7, Q9UQ52, Q96ER9, Q96HA1, H3BR10, P0DMW5, Q96AX1, A8CG34, Q7RTS9, Q16586, Q9Y5W7, Q5JTH9, Q9UKJ5, Q6DT37, Q96PN6, Q5VXU3, Q8TC41, Q5SWX8, Q07812, Q8N138, A2RU48, Q16795, Q06136, Q969S8, Q9Y6U7, Q9H6B9 |
| **d > 35 Å**<br>309 models<br>296 proteins | Q8NFP9, Q86UQ4, P21817, Q9UKL4, P13569, Q2LD37, A6NHN6, Q96HA4, Q8N539, Q07011, Q7Z419, Q9H4F1, E9PIF3, Q6UX15, P0DI80, A6NGB0, Q14318, Q86T13, Q13449, Q13508, Q16620, Q9NY15, O75900, I3L273, P34810, Q9UK23, Q9UMF0, P22748, Q8IZJ0, Q5STR5, Q6N022, Q8N2Q7, P23467, Q9BXS4, O15389, Q8N158, Q9UKZ4, O75581, O14669, Q5VV43, P14415, Q96RD9, P06731, Q7Z407, Q86YD3, Q5VT99, P00387, Q14BN4, E5RHQ5, Q96MH6, O00198, Q71RC9, Q9ULX5, Q8J025, |

| | |
|---|---|
| | Q9NW07, P19256, Q8WUJ3, A0A1B0GUA7, Q9NQC3, Q8IZU9, Q96PQ7, P51636, Q7RTY3, Q9P0K9, Q9Y6N7, O15394, Q14654, O75445, Q4G0I0, P0C7N4, Q9HBL6, B0FP48, O75200, Q16549, A0A087X1L8, P15309, Q8WZ71, Q8NHH1, O14836, Q6UXV1, O95980, Q9UNE0, Q9NPA2, Q6UXN2, Q8WWB7, Q92843, Q5VYJ5, Q9P2E7, Q96CS7, Q15842, Q8NCW0, O15466, P43630, P60608, P43489, Q9NYZ4, A8MVS5, Q96RL6, Q8IWB4, Q96DD7, Q495W5, Q8N326, P10696, P08174, Q14162, Q8NCL9, Q6UW60, P48050, Q13822, Q8N3T6, Q9UND3, A1L1A6, Q8IUK5, O60462, Q53RT3, Q5BVD1, Q86VU5, Q7RTW8, Q16651, O96014, P17927, Q8WYK1, Q9NSU2, Q15768, Q6UXG3, Q8IX05, P26992, Q68D85, Q9Y225, O95210, P56539, P29317, O95944, Q9UNX9, Q4KMG0, O75326, Q9Y334, Q96LZ7, Q86VR7, O14786, Q12913, M0QZC1, Q6UXB8, Q9BZL3, Q9UNG2, P15260, Q3KPI0, Q86V40, O75173, Q15223, P54709, Q96NI6, P98172, Q6DN72, Q9Y644, O95185, Q7L8C5, Q96F15, P35968, P04629, P35052, Q6UXM1, P16109, P29323, Q12836, Q8N387, Q5VUY2, Q9ULH4, Q96L08, O43699, Q9P1W8, P48549, Q9Y6J6, Q9H5Y7, P78508, P51690, Q86SS6, P09912, Q9P2K2, A6NFU0, Q9UH62, P21589, P20138, Q96PD2, Q9H4A9, P40199, P15586, O00763, Q9HD43, Q9NPI9, P22794, P48544, O60235, P16444, Q02763, Q96PQ1, Q8N8G6, Q6ZRH7, Q96Q04, Q14500, F8WFD2, Q96HS1, P20333, P09601, Q5JXA9, Q8WWQ8, Q9HBV2, P07307, Q92806, O60238, Q9BQG1, P52799, P26718, P48051, Q9BXI9, O95274, Q8NCI6, P54753, Q14626, Q8IZA0, Q06643, Q16821, Q16585, Q96FE5, Q8N7C0, Q86SU0, O75144, E9PKD4, Q8N8F7, Q6UXE8, O60449, Q9BYF1, P23470, Q8IW45, P57105, Q5VY43, Q86UE4, Q86YL7, Q9H156, P80370, O43508, Q86T26, P14616, Q9Y6A9, A6NDA9, Q03135, Q9Y6Q6, P0C6S8, Q7Z3B1, A0A2R8Y4M4, Q8N114, B7U540, A0PJX4, P33908, Q7Z553, P48048, Q9NZU1, Q6PL45, H3BTG2, F2Z333, Q9Y336, Q99519, Q16517, Q6PJG9, P55259, Q8WUT4, Q6UWL2, Q6ZVN8, Q8TF66, O00526, A6NFA1, Q6UY18, Q08477, Q9UM22, Q14DG7, Q9NZU0, P0DM63, Q9BT88, P56703, Q9BY79, P41273, P14384, Q7Z692, O60888, Q9BTN0, Q9H665, Q8N7S6, Q7L985, P16581, Q9Y6W8, P07359, Q96EZ4, Q16671, Q6ZMI3, P14679, Q9Y286 |

**Table S2: PFAM entries used to detect transmembrane ABC domains in the 21 proteomes with AF predictions**

| PFAM NAME | PFAM ACC |
|---|---|
| 12TM_1 | PF09847.11 |
| ABC2_membrane | PF01061.26 |
| ABC2_membrane_2 | PF12679.9 |
| ABC2_membrane_3 | PF12698.9 |
| ABC2_membrane_4 | PF12730.9 |
| ABC2_membrane_5[a, b] | PF13346.8 |
| ABC2_membrane_6 | PF06182.13 |
| ABC2_membrane_7 | PF19055.2 |
| ABC_export | PF16962.7 |
| ABC_membrane | PF00664.25 |
| ABC_membrane_2 | PF06472.17 |
| ABC_membrane_3 | PF13748.8 |
| ABC_tran_2 | PF16949.7 |
| BPD_transp_1 | PF00528.24 |
| BPD_transp_2[a,c] | PF02653.18 |
| CbiQ | PF02361.18 |
| CcmB[a,e] | PF03379.15 |
| DUF2705 | PF10920.10 |
| DUF3526 | PF12040.10 |
| DUF3533[a,e] | PF12051.10 |
| DUF6198 | PF19700.1 |
| ECF_trnsprt | PF12822.9 |
| FecCD | PF01032.20 |
| FtsX | PF02687.23 |
| LptF_LptG | PF03739.16 |
| PDR_CDR | PF06422.14 |
| SbmA_BacA[a],[d] | PF05992.14 |
| YitT_membrane[a,b] | PF02588.17 |

[a] PFAM families without associated PDB structures

[b] Our results indicated that YitT is not an ABC fold and incorrectly placed into the ABC-2 PFAM clan

[c] The BPD_transp_2 HMM was identified in bacterial permeases and their AF2-predicted TM structures exhibit BtuC-like fold

[d] AF2 structures of importers with SbmA_BacA HMM hits are Pgp-like

[e] All ABC2_membrane_5, CcmB, and DUF3533 HMM hits include conformations corresponding to the ABCG2-like fold

**Table S3: Opening of zfCFTR structures with kinked TM8 (PDBID: 5W81) and with straight TM8 in MD simulations.**

| system | open frames in each trajectory, # |
|---|---|
| **kinked TM8** | **427 in one and none in the other 21 trajectories*** |
| **straight TM8** | **0, 227, 174, 560, 441, 793**** |

* 22 simulations: 6x100 ns and 16x35ns
* 6 simulations: 6x100 ns
